# Robust quantification of cellular mechanics using optical tweezers

**DOI:** 10.1101/2024.10.13.618126

**Authors:** Wessel S. Rodenburg, Sven F.A. Ebben, Jorine M. Eeftens

## Abstract

Mechanical properties of cells are closely related to function and play a crucial role in many cellular processes, including migration, differentiation, and cell fate determination. Numerous methods have been developed to assess cell mechanics under various conditions, but they often lack accuracy on biologically relevant piconewton-range forces, or have limited control over the applied force. Here, we use optically-trapped polystyrene beads to accurately apply piconewton-range forces to adherent and suspended cells. We precisely apply a constant force to cells by means of a force-feedback system, allowing for quantification of deformation, cell stiffness and creep response from a single measurement. Using drug-induced perturbations of the cytoskeleton, we show that this approach is sensitive to detecting changes in cellular mechanical properties. Collectively, we provide a framework for using optical tweezers to apply highly accurate forces to adherent and suspended cells, and describe straightforward metrics to quantify cellular mechanical properties.

## Introduction

Cells are continuously exposed to mechanical forces *in vivo*. These forces can be generated internally, such as through actomyosin contractions, or can originate from the local environment. For instance, blood and immune cells experience shear stresses in the circulation, migrating cells encounter forces from interactions with the extracellular matrix, and cells within tissues are continuously exposed to compressive stress. The physical and mechanical properties of cells are crucial to sense, resist and respond to such forces, and therefore precisely tuned to align with their function and the local environment^1–5^. The intricate interplay between cell mechanics, mechanosensing, and biological response is vital to many cellular processes, including adhesion, migration, differentiation, and cell fate determination^6–11^. Importantly, alterations in cell mechanics have been associated with aging^12,13^, and disease^14–16^.

The crucial role of cellular mechanics in health and disease has motivated the development of methodologies that apply external forces to cells to measure their response, each with advantages and limitations (recently reviewed in^17^). The most routinely used technique is atomic force microscopy (AFM), in which the tip of a cantilever is used to indent cells. AFM can easily reach high forces but lacks accuracy on smaller, piconewton (pN) range forces – the force range that is key in many biological processes^18,19^. Another commonly applied technique is micropipette aspiration, which deforms cells by applying a local suction pressure. Although relatively easy to implement, micropipette aspiration lacks spatial and temporal resolution, complicating accurate quantification of cellular deformation^20^. Finally, recently developed flow-based techniques, which deform suspended cells through shear fluid forces, enable high-throughput measurements, but have limited options for parallel confocal visualization, and lack control over the applied stress^21^. This results in heterogeneous force application across cells, making robust quantifications challenging.

Most of our current understanding of cell mechanics originates from studies on cells adhered to plastic or glass surfaces. A limitation of this approach is that the properties of the underlying surface can directly influence cell mechanics^2,4,5,8^. Additionally, mechanical perturbations can have profoundly different effects depending on whether cells are adhered to a surface or not^22^. Although technically more challenging, measuring cellular mechanics in the suspended state eliminates any influence from the surface. Therefore, a comprehensive mechanical characterization of cells should include measurements performed on both adherent and suspended cells. Taken together, the ideal technique to study how cells respond to external forces would offer: 1) high accuracy in force quantification; 2) precise control over the applied force and its duration, ensuring all cells undergo the same force; 3) a parallel visual readout (through brightfield and/or confocal microscopy) and 4) the ability to quantify mechanical properties of cells in both adherent and suspended states.

Optical tweezers are exceptional at applying and measuring highly accurate, localized forces. They operate by using a near-infrared focused laser to trap micron-sized beads, with the bead position controllable with nanometer precision. Force in the pN range can be exerted by moving the optical traps towards each other or a sample (squeezing) or moving them away (stretching). While this technique has been widely employed in single-molecule studies^23^ and microrheology^24^, it also holds significant potential as a technique for measuring cellular mechanics^25–27^. However, a detailed description for using optical tweezers for mechanical measurements on cells is currently lacking. In addition, there is no established standard for straightforward quantification of cell mechanics.

Here, we present a robust method to quantify cellular mechanical properties using optical tweezers. We show that mechanical properties of cells can be accurately quantified by applying pN-range forces to adherent and suspended cells using optically-trapped polystyrene beads. To validate the method, we manipulated the cytoskeleton through inhibition of myosin II activity and actin polymerization, showing that these mechanical perturbations significantly soften cells. Our findings demonstrate that optical tweezers are well-suited for precise mechanical measurements on adherent and suspended cells. Finally, we provide an overview of straightforward metrics for quantification of cellular mechanics.

## Results

### Cellular deformation through force application with optically-trapped beads

Optical tweezers can capture and precisely maneuver beads trapped within a focused laser. Force is measured by tracking the displacement of the trapped bead from the center of the optical trap. In this study, we used optically-trapped polystyrene beads (3.15 μm in diameter) to deform adherent and suspended cells. For measuring on adherent cells, a single bead is optically-trapped, positioned at a few μm distance of the side of a cell, and then moved along the lateral direction to deform the cell (**figure 1A**). To visualize the nature and extent of cellular deformation in response to piconewton (pN)-range forces, cells were imaged before and while applying force through a combination of brightfield and confocal microscopy (**figure 1B-C**). Using CellMask, a fluorescent dye that labels the plasma membrane, we clearly observed local indentation of the cell (**figure 1C**). In contrast to adherent cells, measuring on suspended cells requires two optically-trapped beads. One optical trap remains stationary and serves to stabilize the position of the cell, while the other trap moves along the lateral direction (**figure 1D**), which again resulted in small cellular deformations (**figure 1E and supplemental video 1**). We note that the nature of deformation is different between adherent and suspended cells. Whereas adherent cells are locally indented by a single bead, suspended cells are squeezed between two beads, resulting in a global deformation of the cell. Nonetheless, application of pN-range forces deforms cells in the adherent and suspended state.

**Figure 1.**
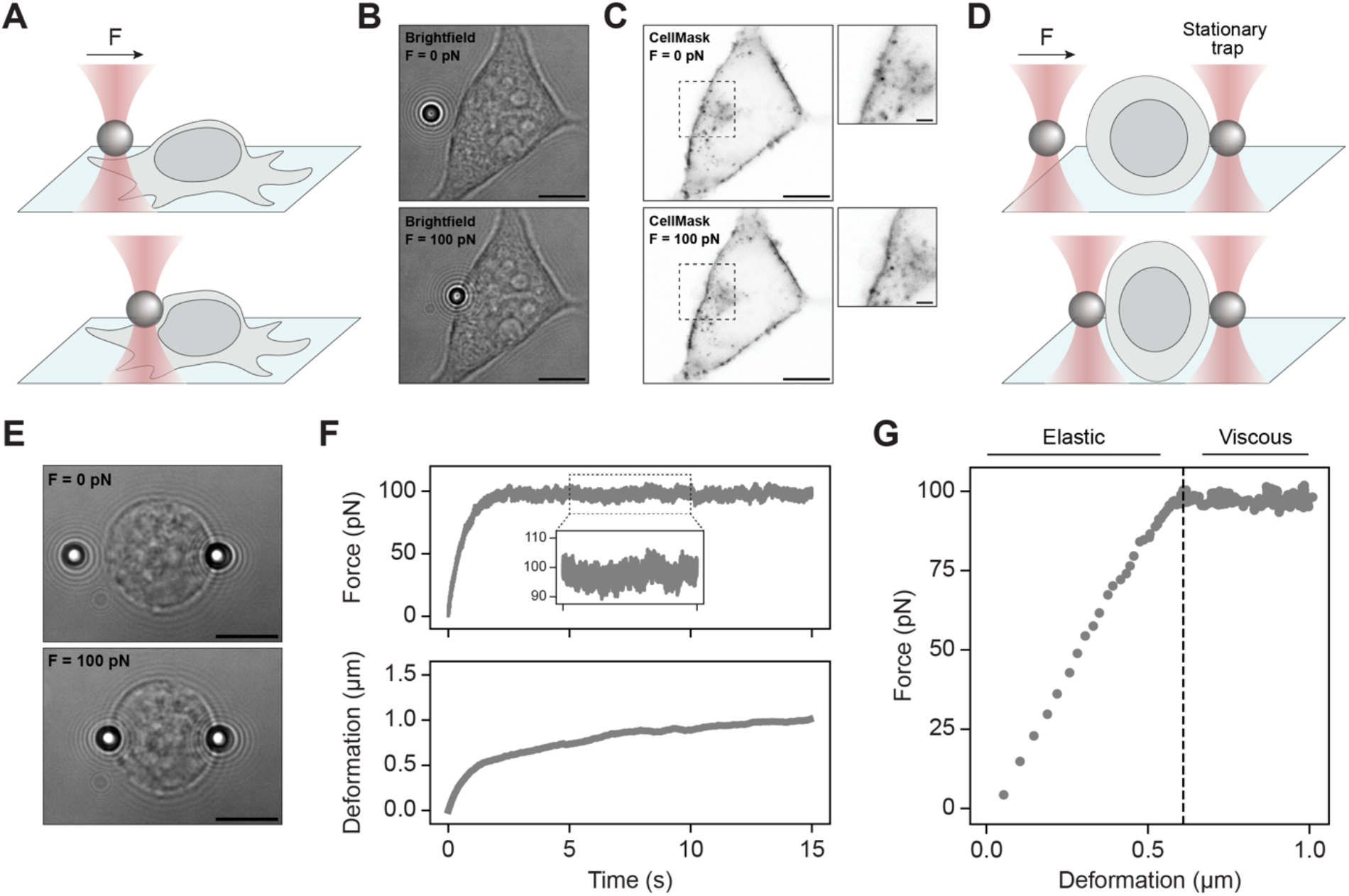
Deformation of adherent and suspended cells using optical tweezers. A) Schematic illustration of an optical tweezers experiment on adherent cells. Cells are seeded on a fibronectin-coated surface. A single bead is optically trapped and moved along the lateral direction to apply force. B) Brightfield images of an adherent cell before force application (top) and during 100 pN force application (bottom). Scale bars, 10 μm. C) Inverted confocal images of the cell membrane (CellMask plasma membrane marker) corresponding with the brightfield images in B. Scale bars, 10 μm. Zoomed-in insets show local deformation. Scale bars, 2 μm. D) Schematic illustration of an optical tweezers experiment on suspended cells. A stationary optically-trapped bead serves to immobilize the cell, while the other is moved along the lateral direction to apply force. E) Brightfield images of a suspended cell before force application (top) and during 100 pN force application (bottom). Scale bars, 10 μm. See also **supplemental video 1**. F) Force (top) and corresponding deformation (bottom) plotted over time for the suspended cell shown in E. A constant load of 100 pN was applied to the cell for ∼10 seconds while the deformation was monitored. G) Force-deformation curve of the suspended cell shown in E. Dashed line indicates the moment the target force of 100 pN is reached.

To take this qualitative observation to a quantifiable metric for cellular deformation in response to force, we track the position of the bead(s) over time. For measurements with adherent cells using one optical trap, the *position* of this single bead over time is sufficient. For measurements with suspended cells using two optical traps, we track the *distance* between the beads over time. As the optical trap approaches the cell, the bead initially moves through liquid and will, at some point, contact the cell. We infer this point of contact from the force-curve (**figure S1A-B**, Materials & Methods). The deformation of the cell at time *t* is subsequently derived by comparing the position of the bead or distance between the beads relative to the point of contact. Besides the precise spatial control over the bead position, another advantage of optical tweezers is the ability to maintain precise control over the applied force, thus ensuring uniformity across measurements. We use a force-feedback system to apply a constant load (or ‘target force’) to cells for at least 10 seconds. This system monitors deviations from the target force at a high frequency and adjusts the optical trap position accordingly, holding the cell in a force-clamp. A typical example of a force-clamp experiment for a suspended cell is shown in **figure 1E-G** and **supplemental video 1**. After initial bead-cell contact, the target force is reached within seconds, and then stably maintained with only minor deviations (**figure 1F**). In the corresponding force-deformation curve, two distinct stages of cellular deformation can be recognized (**figure 1G**). First, while the force is ramping up from 0 to 100 pN, the force response is elastic (force is directly proportional to the deformation). Second, under a constant load of 100 pN, the cell exhibits viscous flow (‘creep’) in which the deformation continues, but at a slower rate (**figure 1F-G**). This characteristic viscoelastic creep behavior is universal among different cell types and has been observed with a variety of techniques^28^. Taken together, the deformation of adherent and suspended cells can be accurately quantified over time while maintaining precise control over the applied force.

### Quantification of cellular mechanical properties

To further standardize our approach for quantifying cellular mechanics, we monitored cellular deformation at two different target forces (50 and 100 pN). The standard deviation from the target force was generally less than 1.5 pN for adherent cells and ∼2-3 pN for suspended cells (**figure S2A**). The deviations are likely slightly higher for suspended cells because these cells, unlike adherent cells, are not fully immobilized. Representative force- and deformation curves for suspended **(figure 2A)** and adherent cells **(figure 2B)** again show a rapid deformation as the force increases, with a creep response once the target force is reached. From these experiments we can now quantify the deformation, which we define as the amount of deformation 10 seconds after reaching the target force. The deformations typically range from 0.2 to 1.5 μm (**figure 2C**), depending on the applied force, with the average deformation increasing as more force is applied (**figure 2C**). As suspended cells are by approximation spherical in shape, we can easily measure the cell diameter and express the deformation as a percentage of the cell diameter (referred to as ‘strain’). The strain of suspended cells typically ranges from 5 to 15%, and increases as more force is applied (**figure S2B**). When we compare the deformation of adherent to suspended cells, we find that adherent cells deform significantly less at the same forces **(figure 2C)**.

**Figure 2.**
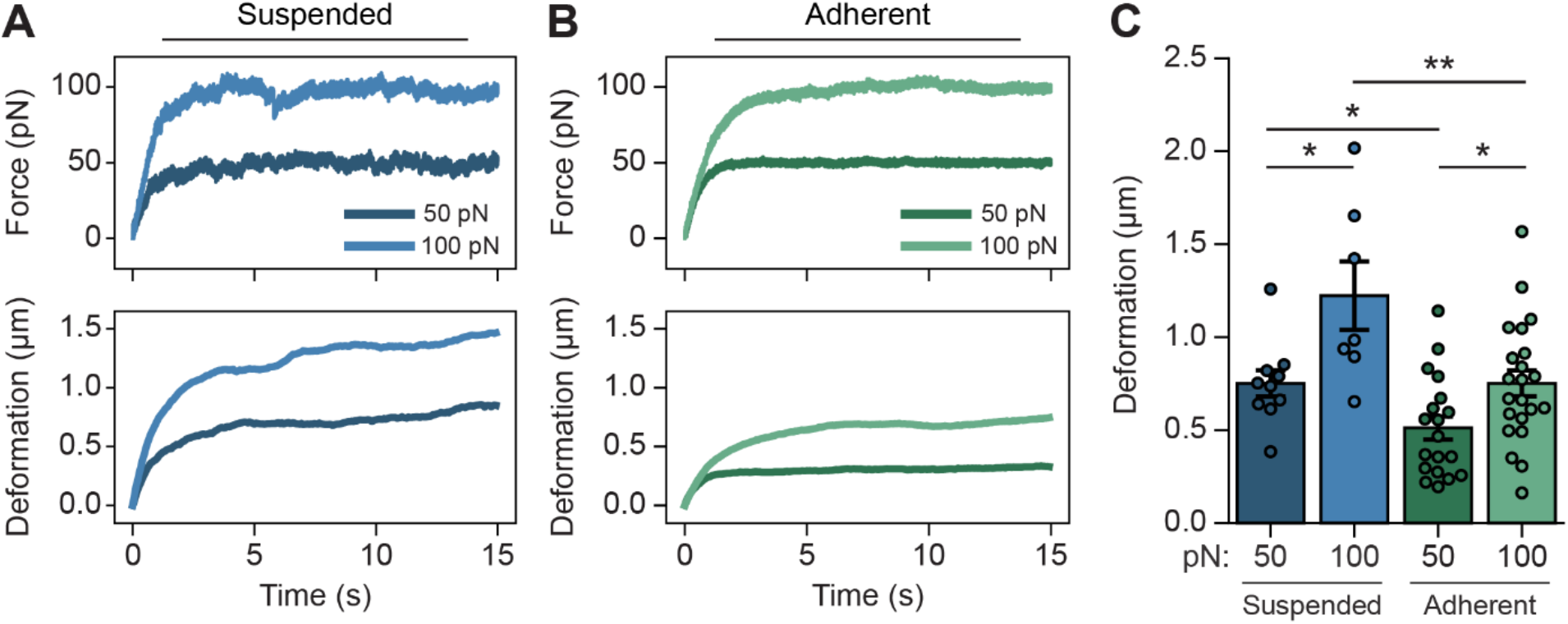
Quantification of cellular deformation. A) Representative force (top) and corresponding deformation (bottom) curves of suspended cells deformed with a target force of 50 pN or 100 pN. B) Representative force (top) and corresponding deformation (bottom) curves of adherent cells deformed with a target force of 50 pN or 100 pN. C) Cellular deformation after 10 seconds of target force application. Data shown as mean ± s.e.m. n = 10, 7, 19 and 22 cells, respectively. *p < 0.05, **p < 0.01, two-sample t-test.

The spring constant is a direct measure of cell stiffness, with a higher spring constant indicating that more force is required to deform the cell. We can calculate the spring constant by linearly fitting the initial elastic force response, where force is directly proportional to the deformation (**figure 3A**). The spring constant is defined as the slope of this fit. When applying a target force of 50 pN, for suspended cells, we find an average spring constant of 99±12pN/μm. For adherent cells, the average spring constant is significantly higher at 267±52 pN/μm. Since we did not yet observe strain stiffening^28^ when applying forces up to 100 pN, the spring constant is expected to be independent from the amount of force applied. Indeed, the average spring constant did not change when increasing the target force from 50 to 100 pN, for both suspended and adherent cells (**figure 3B**). For both target forces, the spring constant for adherent cells is roughly 2-fold lower than for suspended cells **(figure 3B)**.

**Figure 3.**
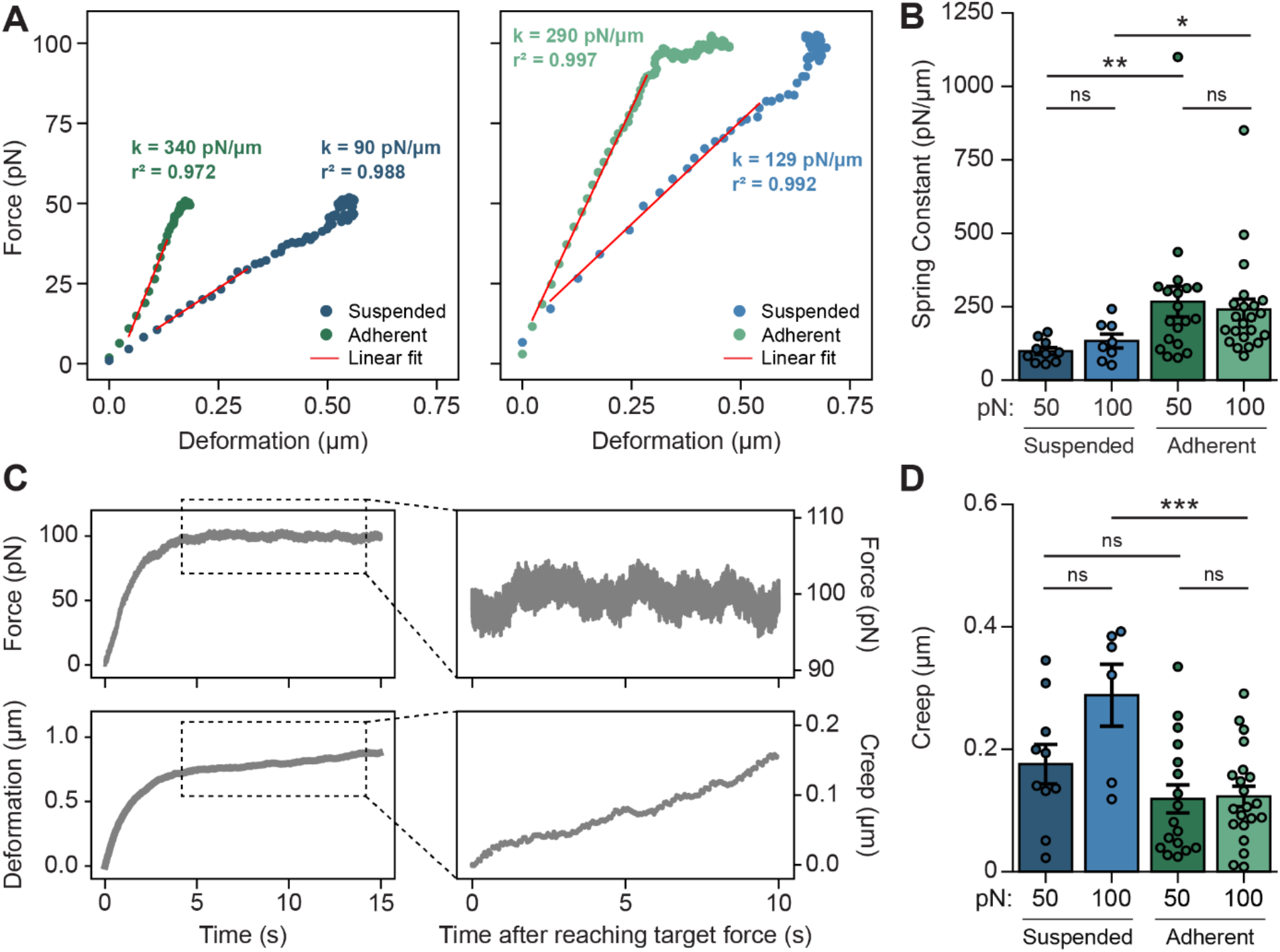
Quantifications of cellular mechanical properties. A) Representative force-deformation curves of a suspended and an adherent cell deformed with a target force of 50 pN (left) and 100 pN (right). Red lines indicate linear fits to calculate the spring constant (k). B) Spring constants calculated from linear fits of individual curves as shown in A. Data shown as mean ± s.e.m. n = 10, 8, 19 and 22 cells, respectively. *p < 0.05, **p < 0.01, Mann-Whitney U test. C) Representative force (top) and corresponding deformation (bottom) curves of an adherent cell deformed with a target force of 100 pN (left), and zoom-in on the creep response (right). The creep response was monitored for 10 seconds after reaching the target force. D) Creep after 10 seconds of target force application. Data shown as mean ± s.e.m. n = 10, 6, 17 and 21 cells, respectively. ***p < 0.001, two-sample t-test.

Next, we focused specifically on the viscous part of the force response: the deformation of cells under a constant force (referred to as ‘creep’) (**figure 3C, inset**). We quantified the amount of creep 10 seconds after reaching the target force, and found that cells continue to deform up to 0.4 μm in this regime (**figure 3D**). Suspended cells generally exhibit more creep deformation than adherent cells, but this difference only reaches significance at 100 pN (**figure 3D**). We noted that the creep response of individual cells is quite heterogeneous, but on average can be described by an inverse exponential decay function: ∆*x* ∗ (1 − *e*^(−*t*/*τ*)^), where ∆x is the creep extent, t is the time, and τ is the relaxation time (**figure S3A-B**). Here, the relaxation time τ is a measure of how quickly the creep deformation levels off under a constant force. The shape of the curves indicates that the creep response has not plateaued yet. Thus, τ offers little insight on this timescale, but could be an insightful metric for longer experiments.

Together, the presented quantifications allow for direct comparison between cells experiencing different forces. In addition, we can compare the stiffness of adherent and suspended cells. The mechanical organization of cells becomes fundamentally different when adherent cells are detached from a substrate, most notably is the absence of stress fibers in suspended cells^22^. In line with these mechanical changes, we found that suspended cells are significantly more deformable at both target forces (**figure 2C**), have a roughly 2-fold lower spring constant (**figure 3B**), and exhibit more creep deformation (**figure 3D**) compared to adherent cells – each suggesting that cells soften when detached from a substrate.

### Validation using pharmacological inhibition of myosin II and actin polymerization

To validate whether our method can successfully quantify changes in cell mechanics due to mechanical perturbations, we applied force to cells treated with inhibitors that are well-known to perturb the cytoskeleton. We first treated cells seeded on a fibronectin-coated surface with 20 μM Blebbistatin: an inhibitor that prevents actomyosin crosslinking by interfering with the ATPase activity of myosin II^29^. Treatment with Blebbistatin has previously been shown to decrease cell stiffness^30–32^. Accordingly, we found that cellular deformation is significantly increased in Blebbistatin treated cells (**figure 4A,C**). Further, the spring constant of cells almost halved in the presence of Blebbistatin (**figure 4B,D**). Thus, adherent cells significantly soften upon inhibition of myosin II activity. Interestingly, we did not find a significant difference in the creep response (**figure 4E, S4A**), suggesting that myosin II inhibition mainly affects the elastic part of the force response. Together, these data thus confirm that mechanical changes on adherent cells can be accurately quantified using optical tweezers.

**Figure 4.**
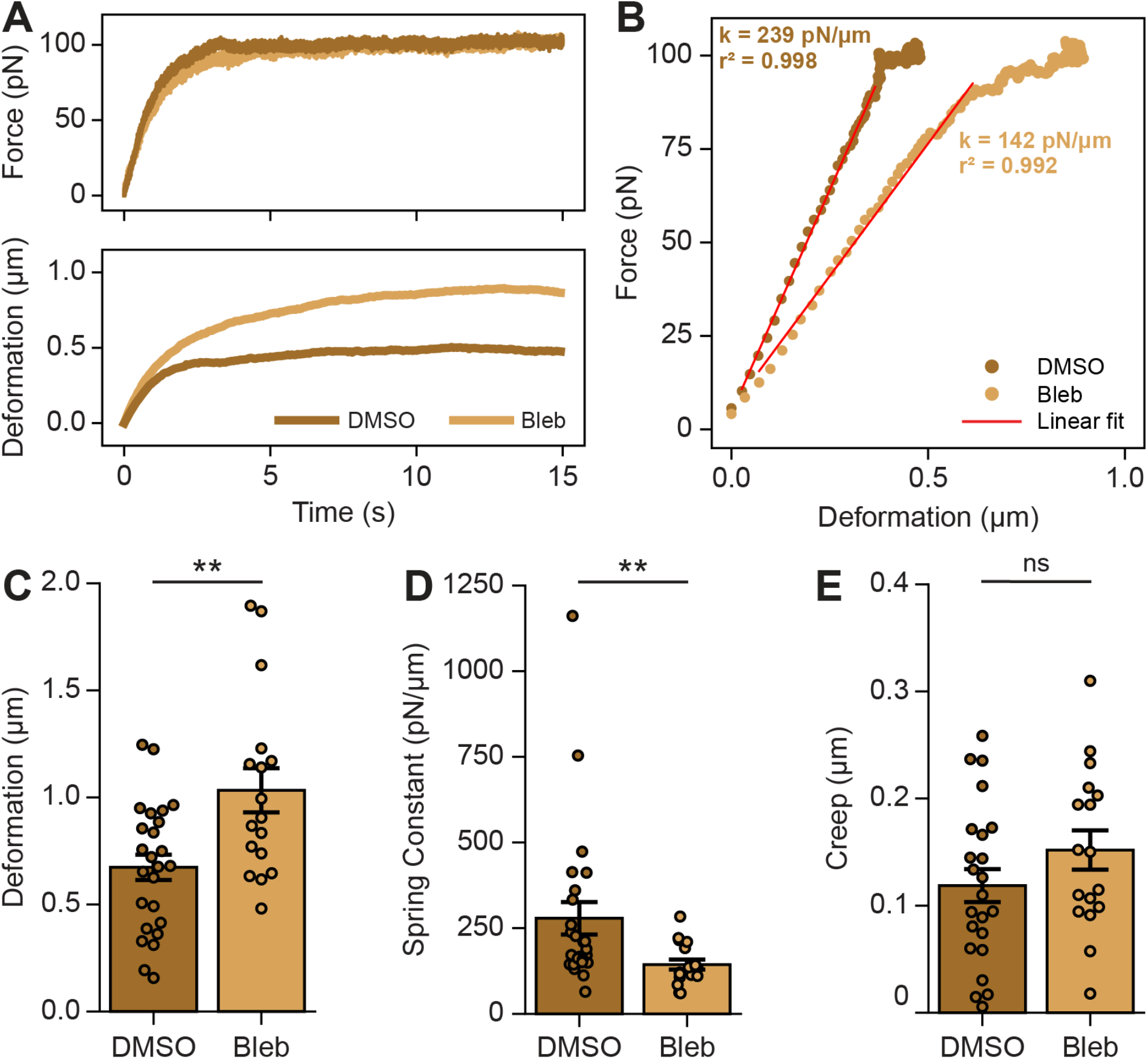
Validation using inhibition of myosin II. A) Representative force (top) and corresponding deformation (bottom) curves of adherent cells treated with DMSO or 20 μM Blebbistatin. B) Representative force-deformation curves of adherent cells treated with DMSO or 20 μM Blebbistatin. Red lines indicate linear fits used to calculate the spring constant (k). C) Cellular deformation after 10 seconds of target force application. Data shown as mean ± s.e.m. n = 25 and 17 cells, respectively. **p < 0.01, two-sample t-test. D) Spring constants calculated from linear fits of individual curves as shown in B. Data shown as mean ± s.e.m. n = 25 and 17 cells, respectively. **p < 0.01, Mann-Whitney U test. E) Creep after 10 seconds of target force application. Data shown as mean ± s.e.m. n = 23 and 17 cells, respectively. p = 0.17, two-sample t-test.

To establish whether our approach is also able to quantify the effect of mechanical perturbations when cells are in suspension, we trypsinized and resuspended cells in medium containing 1 μM Latrunculin-A, an inhibitor that prevents actin polymerization by sequestering monomeric G-actin^33^. When cells are brought in the suspended state, actin forms a thick cortical layer beneath the cell membrane known as the actin cortex, which provides structural and mechanical support^22,34^. Depolymerization of actin filaments using Latrunculin-A is therefore expected to soften cells. Indeed, we found that suspended cells treated with Latrunculin-A are highly deformable, with the average deformation and strain increasing approximately four-fold compared to untreated cells (**figure 5A,C, S5A-B**). Similarly, the spring constant of suspended cells is greatly reduced in the presence of Latrunculin-A (**figure 5B,D**). Finally, Latrunculin-A treated cells exhibited significantly more creep deformation **(figure 5E, S5C)**. These results confirm that the actin cortex of suspended cells is key in providing resistance to external force. Collectively, these data show that optical tweezers can accurately detect changes in mechanical properties upon perturbation when cells are in the suspended state.

**Figure 5.**
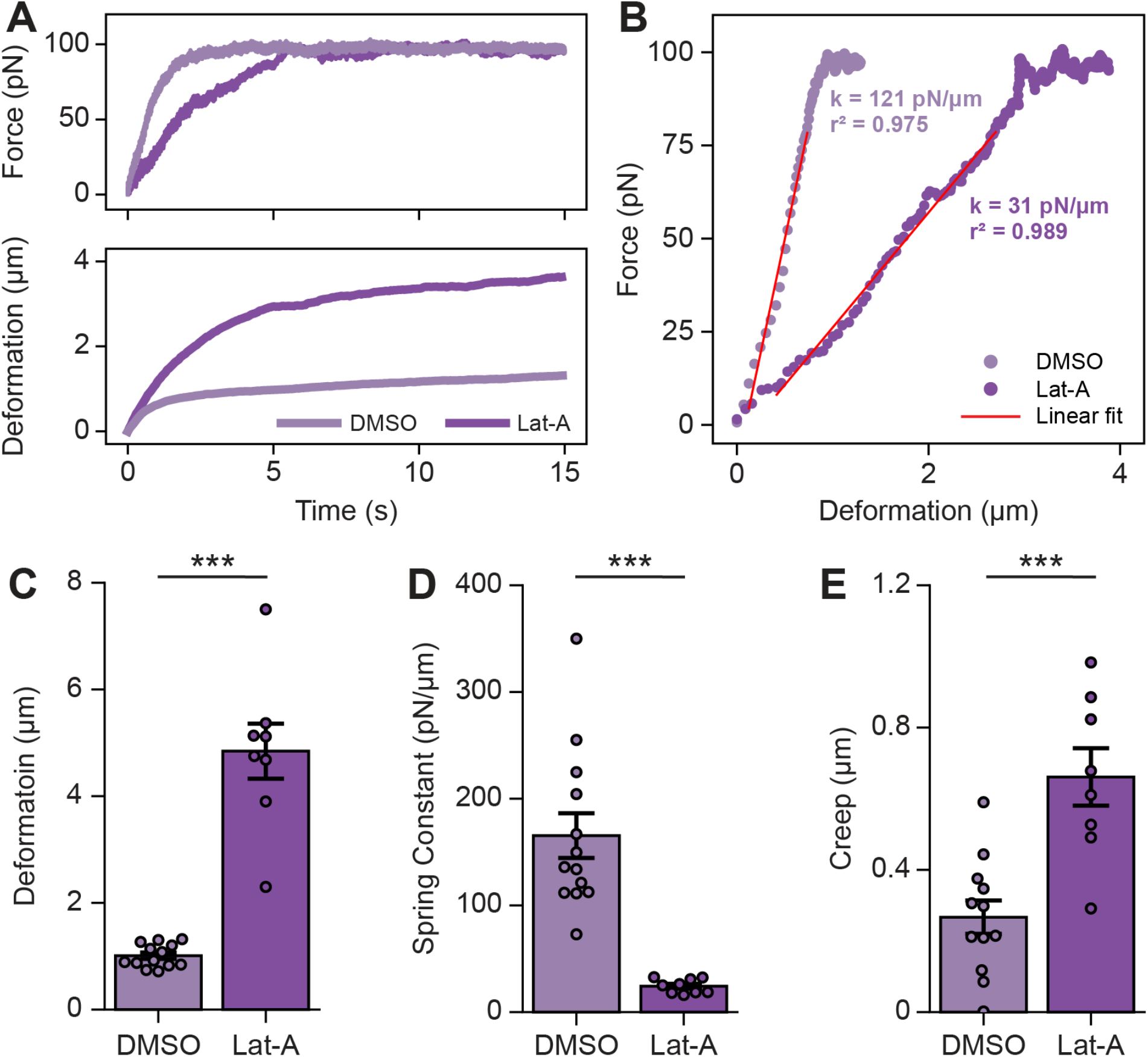
Validation using inhibition of actin polymerization. A) Representative force (top) and corresponding deformation (bottom) curves of suspended cells treated with DMSO or 1 μM Latrunculin-A. B) Representative force-deformation curves of suspended cells treated with DMSO or 1 μM Latrunculin-A. Red lines indicate linear fits used to calculate the spring constant (k). C) Cellular deformation after 10 seconds of target force application. Data shown as mean ± s.e.m. n = 13 and 8 cells, respectively. ***p < 0.001, two-sample t-test. D) Spring constants calculated from linear fits of individual curves as shown in B. Data shown as mean ± s.e.m. n = 13 and 9 cells, respectively. ***p < 0.001, two-sample t-test. E) Creep after 10 seconds of target force application. Data shown as mean ± s.e.m. n = 12 and 8 cells, respectively. ***p < 0.001, two-sample t-test.

## Discussion

Here, we demonstrated the use of optical tweezers for mechanical measurements on adherent and suspended cells. Our approach allows for quantification of cellular deformation, the spring constant and the creep response from a single force-deformation curve. By keeping the optical trap position under control of a force-feedback system, we show that forces can be precisely applied to cells with minimal deviations, thus ensuring uniformity across measurements. Using drug-induced perturbations of the cytoskeleton, we show this approach is sensitive to detecting changes in mechanical properties on both adherent and suspended cells.

Optical tweezers, particularly when equipped with a force-feedback system, allow us to apply the same force to each cell, ensuring experimental robustness. This level of control and accuracy is necessary when making comparisons between conditions, and thus an important advantage of optical tweezers. However, it does come at the expense of a lower force limit, which is typically 200-400 pN. Other techniques such as AFM can reach higher (nN-range) forces, but such forces can induce damage to the cell^18^, and may even deform the underlying substrate^30^. Our results using Blebbistatin and Latrunculin-A show that pN-range forces are sufficient to accurately measure the effect of these perturbations on cellular mechanics. Thus, measurements under controlled but relatively small forces, which are often unattainable with other techniques, are well-suited to accurately measure changes in mechanical properties.

Our current understanding of cell mechanics has mostly been gained from mechanical measurements on adherent cells. We show that optical tweezers can accurately measure on adherent cells, but also on suspended cells, which has several advantages. First, the actin cytoskeleton is heterogeneously organized in adherent cells with stress fibers spanning from one focal adhesion point to another. The measured stiffness of cells can therefore be profoundly different when probed in an area with high or low abundance of stress fibers^35^. Suspended cells have a more homogenously organized cytoskeleton, so quantification of mechanical properties is less dependent on the location of force application. Second, surface modification can directly affect quantification of cell stiffness^4,5,8^, which is ruled out when cells are in the suspended state. Finally, this approach is attractive for mechanical characterization of naturally non-adherent cells such as immune cells, which strongly rely on their mechanical organization for normal functioning^36^.

In conclusion, our findings show that optical tweezers are suitable for robust quantification of cell mechanics. We provide a straightforward method to extract several mechanical properties from a single experiment. The combination with confocal microscopy provides interesting opportunities for visualization of cellular mechanical response to calibrated forces.

## Materials & Methods

### Cell culture

HEK293T cells were cultured in DMEM GlutaMax (Gibco), supplemented with 10% fetal bovine serum (FBS) and 1% penicillin-streptomycin. Cells were grown at 37°C and 5% CO^2^.

### Preparation of cells for optical tweezers experiments

For optical tweezers experiments with adherent cells, μ-Slides (0.4 mm, Ibidi) were pre-coated with 10 μg/ml fibronectin (Sigma-Aldrich, 341631) for 2h at room temperature (RT). Cells were seeded overnight to obtain a confluency of roughly 50%. The medium was then replaced by fresh medium supplemented with 25 mM HEPES (Gibco) containing 3 μl of uncoated polystyrene beads (3.15 μm, 5% w/v, diluted 1:200 in PBS, Spherotech). When combined with confocal imaging as in **figure 1C**, cells were stained with CellMask Green Plasma Membrane Stain (1:1000, ThemoFisher) for 10 min directly prior to starting optical tweezers experiments. Optical tweezers experiments with adherent cells were performed for a maximum of 2-3h at RT.

For experiments with suspended cells, cells grown until roughly 80% confluency were trypsinized, pelleted down, and resuspended in fresh medium supplemented with 25 mM HEPES (Gibco). ∼3000 suspended cells and 3 μl of uncoated polystyrene beads (3.15 μm, 5% w/v, diluted 1:200 in PBS, Spherotech) were added into uncoated μ-Slides (0.4 mm, Ibidi). Cells incubated in the slide for ∼15-30 min prior to starting optical tweezers measurements. Optical tweezers experiments with suspended cells were performed for a maximum of 1-2h at RT.

### Optical trapping and force application

A dual trap optical tweezers setup (C-trap, LUMICKS) equipped with a nanostage was used. Slides were positioned in between a water objective and an oil immersion condenser. IR laser beams (1064 nm) were used for optical trapping of beads. A single bead (for measurements with adherent cells) or two beads (for measurements with suspended cells) were optically-trapped and brought to the appropriate *z*-plane by moving the nanostage along the *z*-axis. For measurements with adherent cells, a single optically-trapped bead is positioned at a few μm distance from the side of a cell (see also **figure 1B**, top image). The optical trap is moved along the lateral (*x*) direction under control of a force-feedback system to apply force to the cell. The force-feedback system quantifies deviations from a pre-defined target force at a rate of 31.3 Hz, and adjusts the optical trap position accordingly with a maximum step size of 50 nm. For measurements with suspended cells, two optically-trapped beads are positioned on opposite sides of a cell along the lateral (*x*) axis, both at a few μm distance of the cell. One bead is manually moved toward the cell along the lateral (*x*) axis until contact is made (see also **figure 1E**, top image). Force is then applied to the cell by moving the other bead along the lateral (*x*) direction under control of a force-feedback system, similar to the measurements with adherent cells. For both adherent and suspended cell measurements, a constant load (i.e., the target force) was applied to the cell for at least 10 seconds. Force data were acquired at a rate of 78125 Hz using back-focal-plane detection. The positions of the beads were acquired at a rate of 15 Hz using bead tracking software (LUMICKS).

### Drug treatments

Latrunculin-A (Sigma, L5163) was diluted to in DMSO and used at a final concentration of 1 μM. Blebbistatin (Sigma, B0560) was diluted in DMSO and used at a final concentration of 20 μM. Cells incubated with the inhibitors (or equivalent amount of DMSO as a control) for ∼15 min prior to starting optical tweezer experiments.

### Data analysis

Data analysis was performed using Pylake Python package (LUMICKS) and custom-made Python scripts. First, the starting point of deformation (i.e., point of contact between the bead and the cell) is calculated for each measurement. We defined the point of contact as the *bead position* (for adherent cells) or the *distance between the beads* (for suspended cells) when force first reaches a value below 0 pN (starting from the maximum force) (see also **figure S1A-B**). The deformation of the cell is calculated over time by comparing the position of the bead or the distance between the beads relative to the point of contact. Mechanical properties are extracted from the resulting force-deformation curves. The deformation was defined as the amount of deformation 10 seconds after reaching the target force. The spring constant was calculated as the slope of the linear part of the force-deformation curve through linear regression analysis. For suspended cells, we additionally quantified the cell diameter to derive the strain – that is, the ratio of deformation over the cell diameter. The cell diameter was quantified as the distance between the center of the two beads at the point of contact, minus the bead diameter. The creep deformation was defined as the additional deformation measured 10 seconds after reaching the target force. The average creep response was fitted to (1):

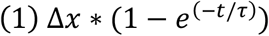

from which best fit values of Δx (creep extent) and τ (relaxation time) were derived. In 7 out of 121 measurements, the cell exhibited negative creep (i.e., the bead being pushed back by the cell), which were excluded from the analysis of the creep response.

### Statistical analysis

Datasets were tested for normality using Shapiro-Wilk test (α = 0.05). Two-sample *t*-tests were used for comparing normally distributed data. Non-normally distributed data were compared using Mann-Whitney U test. In both cases, *p*-values lower than 0.05 were considered statistically significant. Data are presented as mean ± s.e.m.

## Acknowledgements

We thank the Radboud UMC Technology Center Microscopy for use of their microscopy facilities, and Marieke Willemse and Koen van den Dries for generously sharing reagents.

## Supplemental Figures and Video

**Figure S1.**
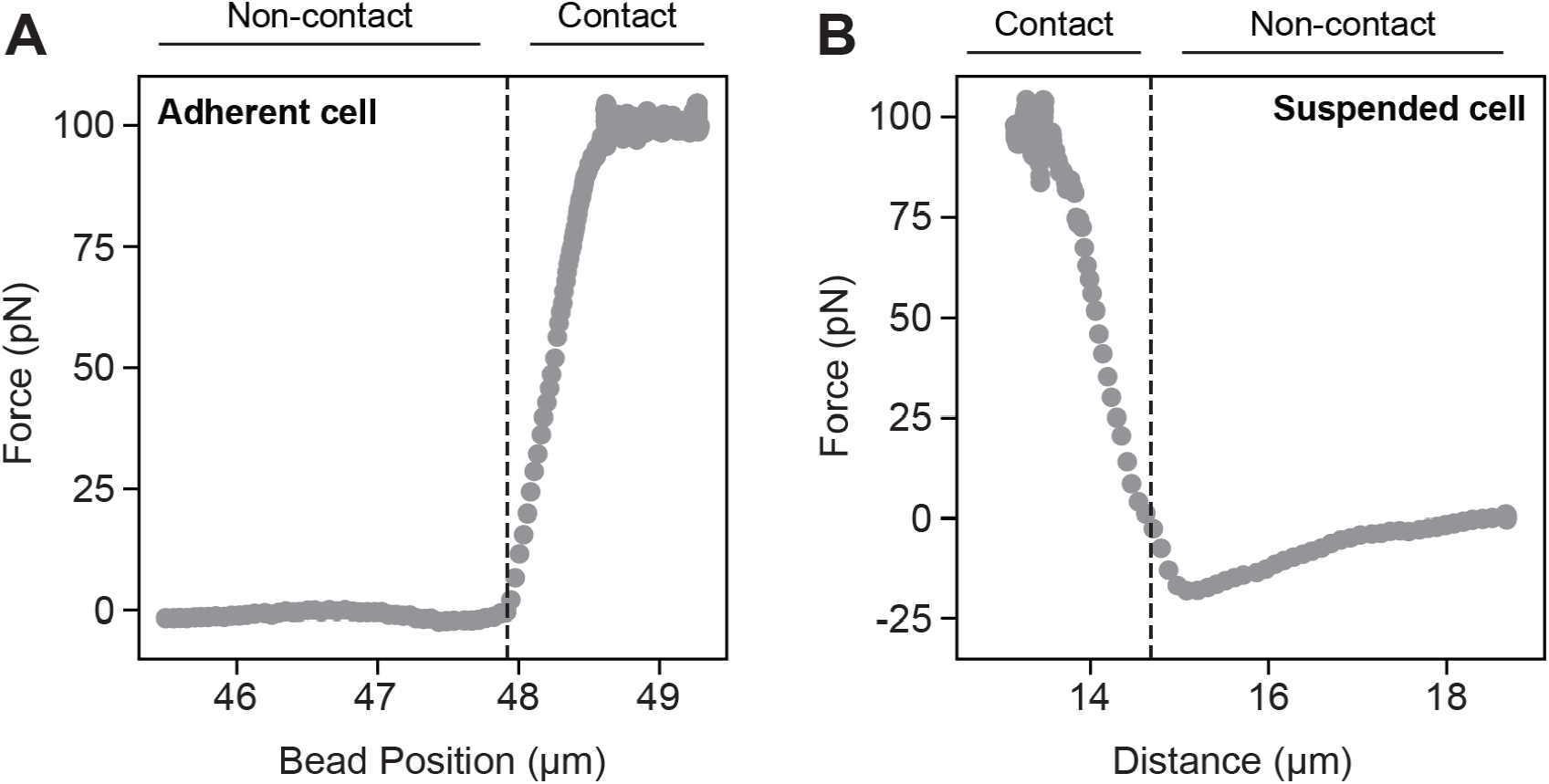
Defining the point of bead-cell contact. A) Representative force-bead position plot for an adherent cell deformed with a target force of 100 pN. Dashed line indicates the point of contact (i.e., the start of deformation). B) Representative force-distance plot for a suspended cell deformed with a target force of 100 pN. Dashed line indicates the point of contact (i.e., the start of deformation).

**Figure S2.**
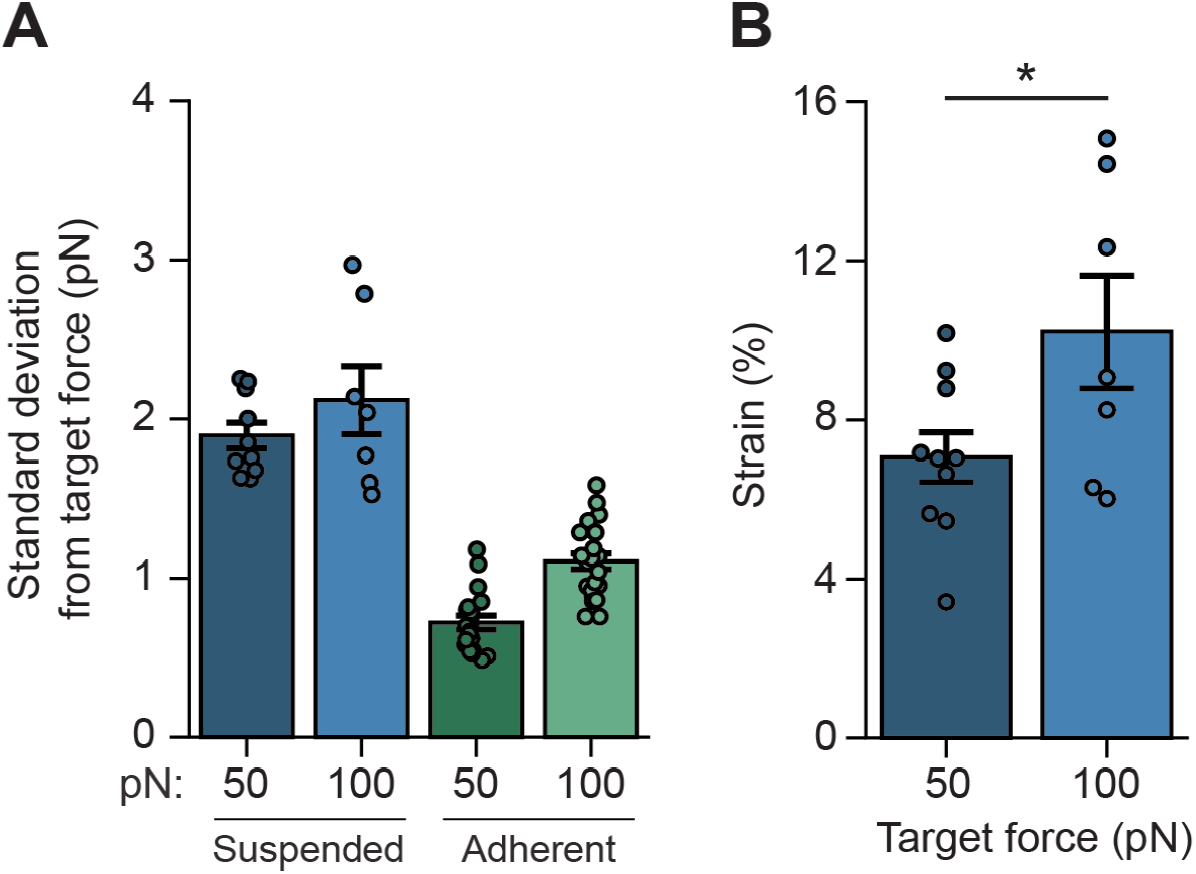
Validating cellular deformation at constant force. A) Standard deviation from the target force for the measurements used in **figure 2**. Data shown as mean ± s.e.m. B) Strain of suspended cells after 10 seconds of target force application. Strain is calculated as the ratio of deformation over the cell diameter, expressed as a percentage. Data shown as mean ± s.e.m. n = 10 and 7 cells, respectively. *p < 0.05, two-sample t-test.

**Figure S3.**
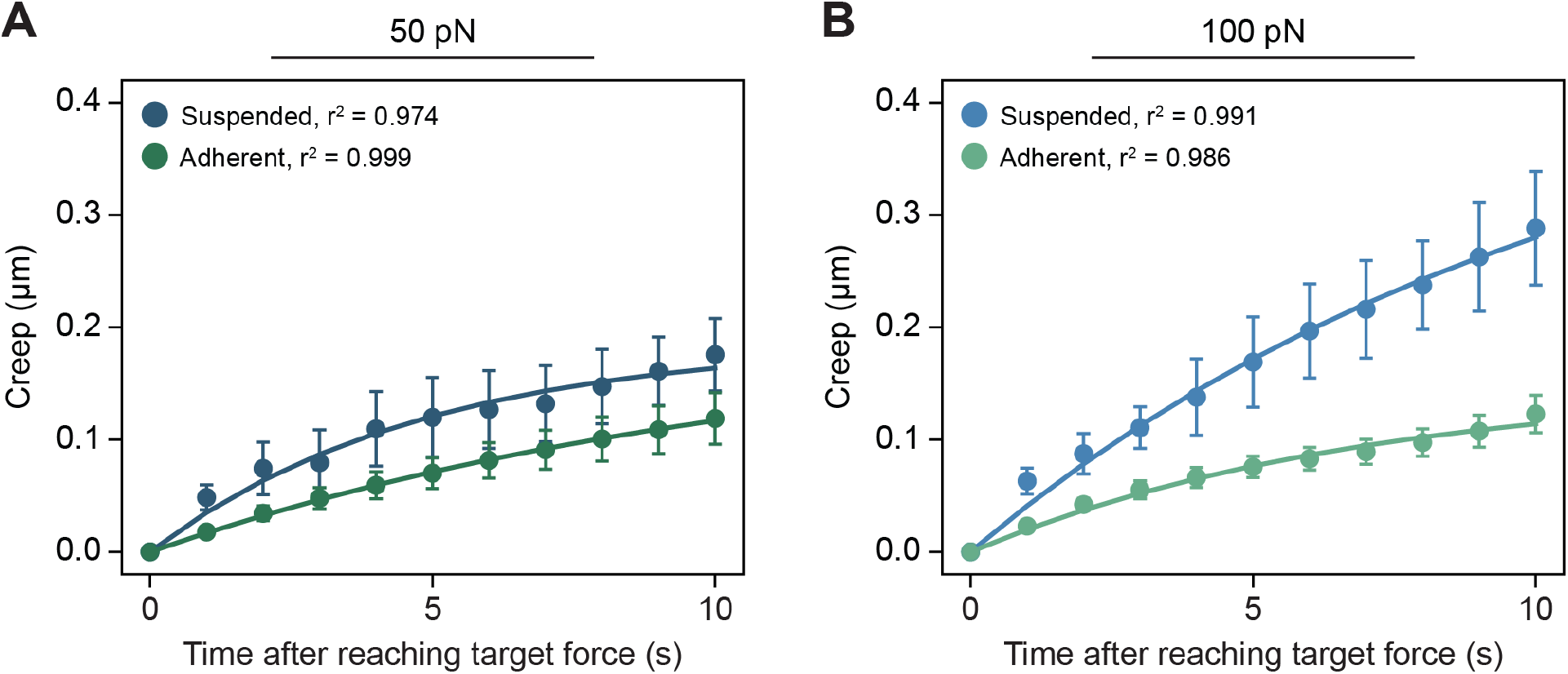
Average creep response of suspended and adherent cells. A) Average creep deformation curve of suspended and adherent cells under a constant load of 50 pN. The creep response was monitored for 10 seconds after reaching the target force. Data shown as mean ± s.e.m. Data is fitted by 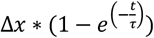. Best fit values are ∆*x* = 0.19 μm and *τ* = 4.8 s for suspended cells, and ∆*x* = 0.20 μm and *τ* = 11.2 s for adherent cells. B) Average creep deformation curve of suspended and adherent cells under a constant load of 100 pN. The creep response was monitored for 10 seconds after reaching the target force. Data shown as mean ± s.e.m. Data is fitted by 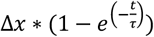. Best fit values are ∆*x* = 0.46 μm and *τ* = 10.8 s for suspended cells, and ∆*x* = 0.15 μm and *τ* = 7.0 s for adherent cells.

**Figure S4.**
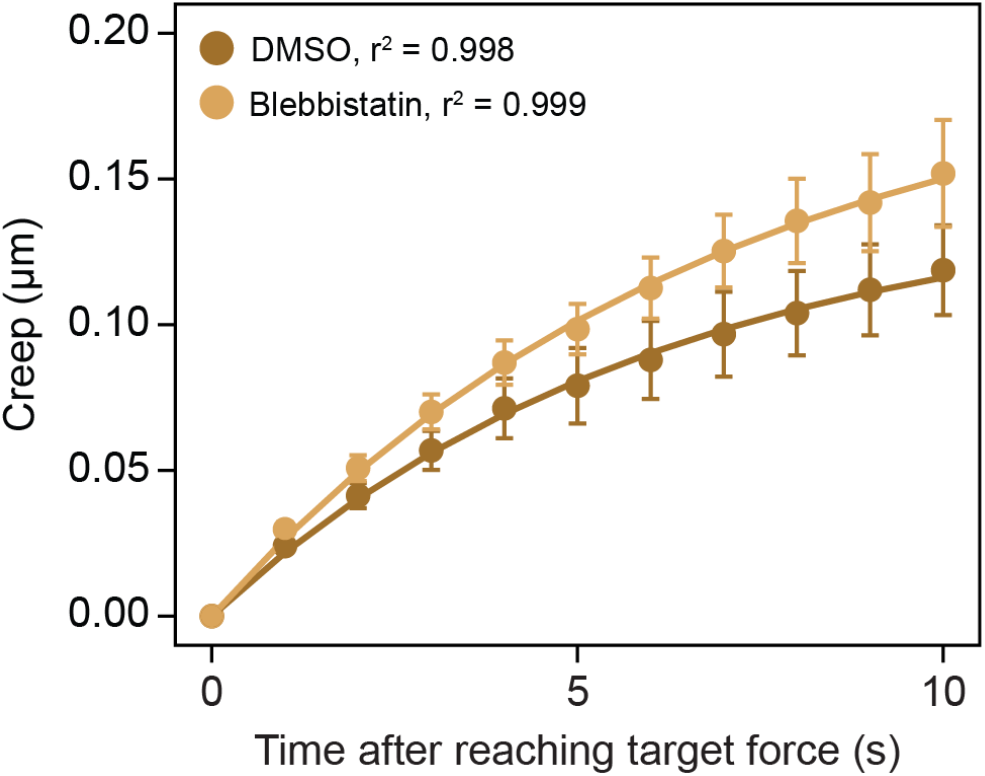
Average creep response of Blebbistatin-treated cells compared to controls. Average creep deformation of adherent cells treated with DMSO or 20 μM Blebbistatin under a constant load of 100 pN. The creep response was monitored for 10 seconds after reaching the target force. Data shown as mean ± s.e.m. Data is fitted by 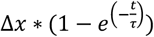. Best fit values are ∆*x* = 0.14 μm and *τ* = 6.1 s for DMSO, and ∆*x* = 0.20 μm and *τ* = 6.8 s for Blebbistatin.

**Figure S5.**
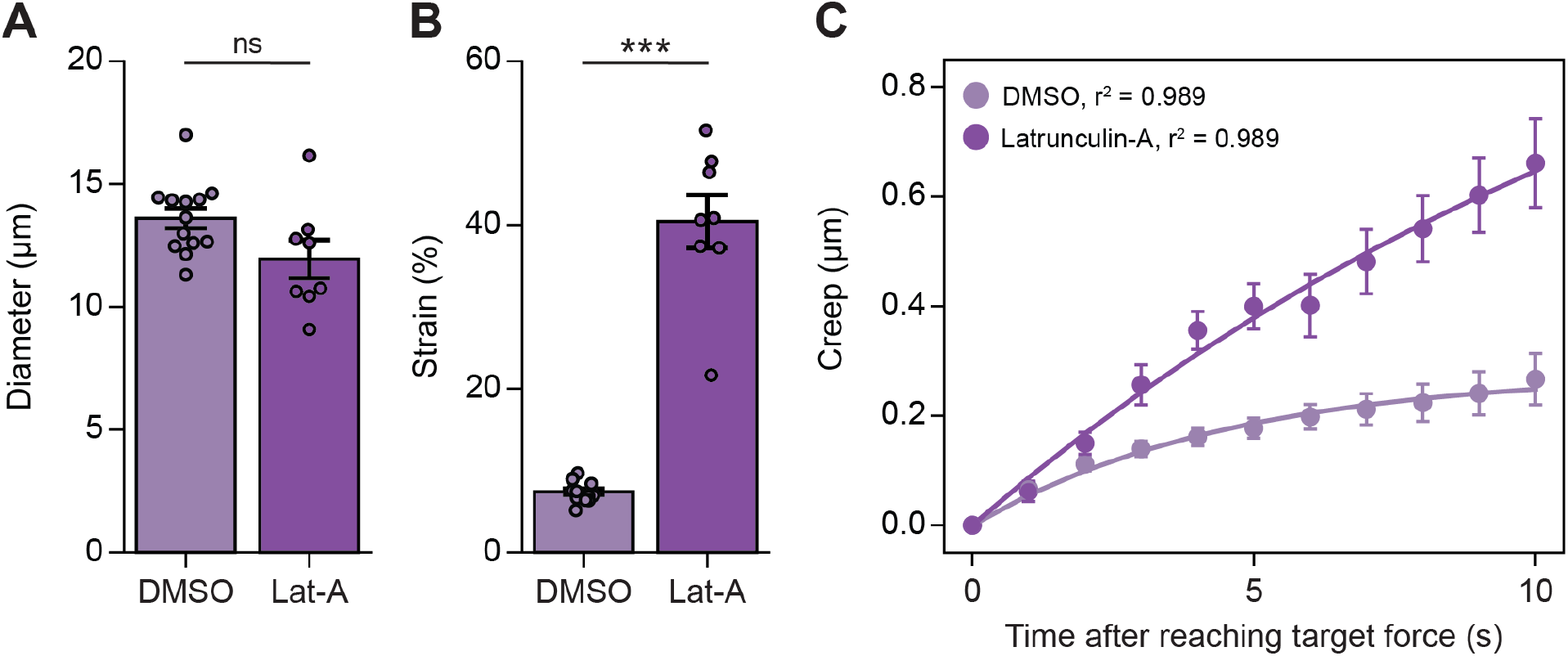
Validating mechanical diHerences between Latrunculin-A treated cells and controls through additional quantifications. A) Diameter of suspended cells treated with DMSO or 1 μM Latrunculin-A. Data shown as mean ± s.e.m. n = 13 and 8 cells, respectively. p = 0.05, two-sample t-test. B) Strain of suspended cells treated with DMSO or 1 μM Latrunculin-A. Strain is calculated as the ratio of deformation over the cell diameter, expressed as a percentage. n = 13 and 8 cells, respectively. ***p < 0.001, two-sample t-test. C) Average creep deformation of suspended cells treated with DMSO or 1 μM Latrunculin-A under a constant load of 100 pN. The creep response was monitored for 10 seconds after reaching the target force. Data shown as mean ± s.e.m. Data is fitted by 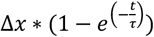. Best fit values are ∆*x* = 0.28 μm and *τ* = 4.5 s for DMSO, and ∆*x* = 1.29 μm and *τ* = 14.4 s for Latrunculin-A.

---

**Supplementary video 1: Force-clamp measurement on a suspended cell**.

Example of a force-clamp experiment for a suspended cell, related to figures 1E-G. One bead remains stationary to stabilize the position of the cell, while the other applies force. Video is shown in real-time with a framerate of 15 Hz. Scale bar, 10 μm.

## References

1. Martino, F., Perestrelo, A. R., Vinarský, V., Pagliari, S. & Forte, G. Cellular Mechanotransduction: From Tension to Function. Front. Physiol. 9, 824 (2018).

2. Discher, D. E., Janmey, P. & Wang, Y. Tissue Cells Feel and Respond to the Stiffness of Their Substrate. Science 310, 1139–1143 (2005).

3. Mathur, A. B., Collinsworth, A. M., Reichert, W. M., Kraus, W. E. & Truskey, G. A. Endothelial, cardiac muscle and skeletal muscle exhibit different viscous and elastic properties as determined by atomic force microscopy. Journal of Biomechanics 34, 1545–1553 (2001).

4. Solon, J., Levental, I., Sengupta, K., Georges, P. C. & Janmey, P. A. Fibroblast Adaptation and StiOness Matching to Soft Elastic Substrates. Biophysical Journal 93, 4453–4461 (2007).

5. Janmey, P. A., Fletcher, D. A. & Reinhart-King, C. A. StiOness Sensing by Cells. Physiological Reviews 100, 695–724 (2020).

6. Lautenschläger, F. et al. The regulatory role of cell mechanics for migration of differentiating myeloid cells. Proc. Natl. Acad. Sci. U.S.A. 106, 15696–15701 (2009).

7. Dupont, S. & Wickström, S. A. Mechanical regulation of chromatin and transcription. Nat Rev Genet 23, 624–643 (2022).

8. Engler, A. J., Sen, S., Sweeney, H. L. & Discher, D. E. Matrix Elasticity Directs Stem Cell Lineage Specification. Cell 126, 677–689 (2006).

9. Vining, K. H. & Mooney, D. J. Mechanical forces direct stem cell behaviour in development and regeneration. Nat Rev Mol Cell Biol 18, 728–742 (2017).

10. Shiraishi, K. et al. Biophysical forces mediated by respiration maintain lung alveolar epithelial cell fate. Cell 186, 1478-1492.e15 (2023).

11. Sun, Y. et al. Mechanics Regulates Fate Decisions of Human Embryonic Stem Cells. PLoS ONE 7, e37178 (2012).

12. Phillip, J. M., Aifuwa, I., Walston, J. & Wirtz, D. The Mechanobiology of Aging. Annu. Rev. Biomed. Eng. 17, 113–141 (2015).

13. Bajpai, A., Li, R. & Chen, W. The cellular mechanobiology of aging: from biology to mechanics. Annals of the New York Academy of Sciences 1491, 3–24 (2021).

14. Wirtz, D., Konstantopoulos, K. & Searson, P. C. The physics of cancer: the role of physical interactions and mechanical forces in metastasis. Nat Rev Cancer 11, 512–522 (2011).

15. Gensbittel, V. et al. Mechanical Adaptability of Tumor Cells in Metastasis. Developmental Cell 56, 164–179 (2021).

16. Zuela-Sopilniak, N. & Lammerding, J. Can’t handle the stress? Mechanobiology and disease. Trends in Molecular Medicine 28, 710–725 (2022).

17. Hobson, C. M., Falvo, M. R. & Superfine, R. A survey of physical methods for studying nuclear mechanics and mechanobiology. APL Bioengineering 5, 041508 (2021).

18. Haase, K. & Pelling, A. E. Investigating cell mechanics with atomic force microscopy. J. R. Soc. Interface. 12, 20140970 (2015).

19. Yusko, E. C. & Asbury, C. L. Force is a signal that cells cannot ignore. MBoC 25, 3717–3725 (2014).

20. Hochmuth, R. M. Micropipette aspiration of living cells. Journal of Biomechanics 33, 15–22 (2000).

21. Urbanska, M. et al. A comparison of microfluidic methods for high-throughput cell deformability measurements. Nat Methods 17, 587–593 (2020).

22. Chan, C. J. et al. Myosin II Activity Softens Cells in Suspension. Biophysical Journal 108, 1856–1869 (2015).

23. Bustamante, C. J., Chemla, Y. R., Liu, S. & Wang, M. D. Optical tweezers in single-molecule biophysics. Nat Rev Methods Primers 1, 25 (2021).

24. Robertson-Anderson, R. M. Optical Tweezers Microrheology: From the Basics to Advanced Techniques and Applications. ACS Macro Lett. 7, 968–975 (2018).

25. Dao, M., Lim, C. T. & Suresh, S. Mechanics of the human red blood cell deformed by optical tweezers. Journal of the Mechanics and Physics of Solids 51, 2259–2280 (2003).

26. Zhang, H. & Liu, K.-K. Optical tweezers for single cells. J R Soc Interface 5, 671–690 (2008).

27. Català-Castro, F., Schäffer, E. & Krieg, M. Exploring cell and tissue mechanics with optical tweezers. Journal of Cell Science 135, jcs259355 (2022).

28. Moeendarbary, E. & Harris, A. R. Cell mechanics: principles, practices, and prospects. WIREs Mechanisms of Disease 6, 371–388 (2014).

29. Kovács, M., Tóth, J., Hetényi, C., Málnási-Csizmadia, A. & Sellers, J. R. Mechanism of Blebbistatin Inhibition of Myosin II. Journal of Biological Chemistry 279, 35557–35563 (2004).

30. Rheinlaender, J. et al. Cortical cell stiffness is independent of substrate mechanics. Nat. Mater. 19, 1019–1025 (2020).

31. Moeendarbary, E. et al. The cytoplasm of living cells behaves as a poroelastic material. Nature Mater 12, 253–261 (2013).

32. Martens, J. C. & Radmacher, M. Softening of the actin cytoskeleton by inhibition of myosin II. Pflugers Arch - Eur J Physiol 456, 95–100 (2008).

33. Coué, M., Brenner, S. L., Spector, I. & Korn, E. D. Inhibition of actin polymerization by latrunculin A. FEBS Letters 213, 316–318 (1987).

34. Flormann, D. A. D. et al. The structure and mechanics of the cell cortex depend on the location and adhesion state. Proc. Natl. Acad. Sci. U.S.A. 121, e2320372121 (2024).

35. Maloney, J. M. et al. Mesenchymal Stem Cell Mechanics from the Attached to the Suspended State. Biophysical Journal 99, 2479–2487 (2010).

36. Huse, M. Mechanical forces in the immune system. Nat Rev Immunol 17, 679–690 (2017).

